# Quantification of beta cell carrying capacity in prediabetes

**DOI:** 10.1101/2024.01.31.578144

**Authors:** Aurore Woller, Yuval Tamir, Alon Bar, Avi Mayo, Michal Rein, Anastasia Godneva, Netta Mendelson Cohen, Eran Segal, Yoel Toledano, Smadar Shilo, Didier Gonze, Uri Alon

## Abstract

Prediabetes, a subclinical state of high glucose, carries a risk of transition to diabetes. One cause of prediabetes is insulin resistance, which impairs the ability of insulin to control blood glucose. However, many individuals with high insulin resistance retain normal glucose due to compensation by enhanced insulin secretion by beta cells. Individuals seem to differ in their maximum compensation level, termed beta cell carrying capacity, such that low carrying capacity is associated with a higher risk of prediabetes and diabetes. Carrying capacity has not been quantified using a mathematical model and cannot be estimated directly from measured glucose and insulin levels in patients, unlike insulin resistance and beta cell function which can be estimated using HOMA-IR and HOMA-B formula.

Here we present a mathematical model of beta cell compensation and carrying capacity, and develop a new formula called HOMA-C to estimate it from glucose and insulin measurements. HOMA-C estimates the maximal potential beta cell function of an individual, rather than the current beta cell function. We test this approach using longitudinal cohorts of prediabetic people, finding 10-fold variation in carrying capacity. Low carrying capacity is associated with higher risk of transitioning to diabetes. We estimate the timescales of beta cell compensation and insulin resistance using large datasets, showing that, unlike previous mathematical models, the new model can explain the slow rise in glucose over decades. Our mathematical understanding of beta cell carrying capacity may help to assess the risk of prediabetes in each individual.

## Introduction

Prediabetes is defined as a state in which fasting glucose is between 5.6mM and 6.9 mM (1,2) with global prevalence estimated at 10-30% (3). Additional criteria for diagnosis include OGTT and HbA1C criteria (2). Prediabetes is a subclinical state without symptoms. However, prediabetes can transition to type-2 diabetes, in which glucose exceeds 7mM, with a yearly conversion rate of 5-10% (1). Since type-2 diabetes is associated with morbidity and mortality (2), it is important to understand the origin of prediabetes and to determine which individuals are at risk to transition to diabetes.

One of the main causes of prediabetes is rising insulin resistance (4,5). In insulin resistance, each unit of insulin is less effective in removing glucose from the blood. Insulin resistance can change rapidly within hours to days in conditions such as inflammation, pregnancy and acute stress. In these situations, insulin resistance is modulated by hormones and cytokines, and is thought to be a physiological knob that allocates glucose appropriately (6). Insulin resistance also rises in obesity due to inflammation and fat accumulation in muscles and liver. Insulin resistance and beta cell function can be estimated for research purposes from glucose and insulin blood tests using mathematical-model-based formula such as HOMA-IR and HOMA-B and their more recent improvements (7,8).

Importantly, however, chronically high insulin resistance in many situations does not lead to abnormal glucose levels. This includes most people with obesity, who have high insulin resistance but normal glucose levels (5). The reason for this is that beta cell function can rise to compensate for insulin resistance (5,9,10,11). This increase in function is due to increased insulin secretion per unit beta cell biomass, and increased total beta cell biomass due to hypertrophy. The total ability of beta cells to secrete insulin is called the beta cell functional mass. As beta cell functional mass rises, more insulin is secreted, exactly compensating for insulin resistance (12). People with obesity indeed have increased beta cell functional mass (13) and increased insulin levels (14).

This raises the question of how prediabetes might occur. Why is compensation not enough to counteract insulin resistance and maintain normal glucose levels and thus prevent prediabetes? One current explanation is metabolic variation between individuals. More specifically, individuals can differ in the maximal compensation enabled by beta cells (maximal potential insulin secretion capacity), which can be called the beta cell carrying capacity (15). Variations in carrying capacity may be due to differences in the number of beta cells reached during development, as found in postmortem studies (16). Carrying capacity is thus suggested to be predictive of the risk for prediabetes and diabetes. Individuals with low carrying capacity would be at risk because a given level of insulin resistance would cause higher glucose than in those with high carrying capacity.

However, the concept of beta cell carrying capacity has not been made quantitative using a mathematical approach. Unlike insulin resistance and beta cell function, which have well characterized mathematical models leading to useful research formulas like HOMA-IR and HOMA-B, there are no such mathematical models for carrying capacity, and there is no analogous HOMA-like formula to estimate the carrying capacity in patients. Therefore, we lack quantitative measurements and mathematical characterization of a potential mechanistic factor that can help to understand some of the variation in prediabetes between individuals.

Here we present a mathematical model of beta cell function and glucose dynamics which includes a carrying capacity for beta cell functional mass. We use the model to develop a HOMA-like formula for carrying capacity based on insulin and glucose blood tests. We call this HOMA-C. It aims to estimate the maximal potential beta cell function achievable by an individual with prediabetes. We test this model using longitudinal datasets of unmedicated patients with prediabetes and newly diagnosed diabetes (17,18). We find that carrying capacity varies by a factor of 10 between individuals and does not vary strongly with age. We also find that Iow carrying capacity is associated with a high risk of diabetes. Finally, we calibrate the timescale of compensation in the model using large datasets and discuss diabetes subtypes in light of beta cell carrying capacity.

## Results

### Mathematical model with beta cell carrying capacity can explain prediabetes

To develop a model for beta cell carrying capacity, we use as a basis the classical mathematical model for diabetes, exemplified by the HOMA model (7). We add to the classical model an equation for beta cell growth, based on the model of Topp et al (9). The crucial new feature in our model that goes beyond existing models is a limit to the total beta cell function, the *beta cell carrying capacity*.

This carrying capacity in humans is based on two physiological limits. First, there is a maximum secretion rate per unit biomass that a beta cell is capable of. This maximum is due to the fact that there is only a finite amount of protein machinery that can fit into a unit of biomass (19). Second, there is a maximum to the total biomass summed over all beta cells in the body. In humans, beta cells do not proliferate significantly after childhood (20). They can increase their size (hypertrophy) to compensate for high glucose levels. However, cells cannot increase their size indefinitely. Recent data suggests that cells that grow to about twice their normal volume begin to resemble senescent cells and secrete proinflammatory signals (21). Human data suggest that total beta cell mass grows by a factor of less than 2 in obesity (22,23,13).

The carrying capacity is thus defined as the product of the maximal insulin secretion per unit biomass times the maximal total biomass of all beta cells in an individual. The carrying capacity mathematical model includes a term that forces beta cell functional mass growth to become zero when mass *B* reaches a carrying capacity *C*.

We begin with a classical model without carrying capacity that accounts for beta cell compensation, a simplified version of the model developed by Topp et al (9). The model has three equations:

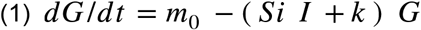

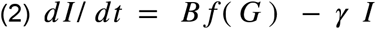

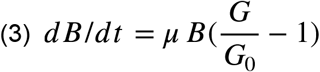

where *m*_0_ is a glucose source term that accounts for liver glucose production plus meal intake, *Si* is insulin sensitivity and *k* is the insulin-independent glucose removal rate. Insulin is secreted by beta cells with functional mass *B* (in units of insulin concentration per unit time) and removed at rate *γ*. The regulation function of beta cell insulin secretion by glucose is a sigmoidal function 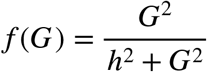 with halfway point *h* (24). The beta cell functional mass changes with growth rate to achieve the glucose set point *G*_0_.

This model accounts for compensation on the timescale of weeks, as beta cell functional mass grows to buffer changes in parameters such as insulin sensitivity *Si* (10).

To develop a model for the carrying capacity, we add to the beta cell growth equation a limit for growth that is inspired by ecological carrying capacity:

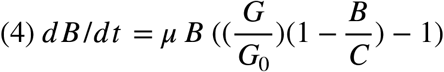

As beta cell functional mass approaches C, its growth rate slows down, according to a linear term 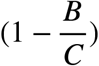 adapted from carrying capacity in ecology and in cellular replication (25,26,27). In this model, there is sufficient compensation when *B* is much smaller than the carrying capacity, where we can approximate 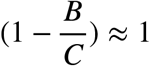 and return to the Topp model. However, when *B* approaches carrying capacity *C*, compensation begins to break down. Insulin production becomes less than needed for sufficient compensation. As a result, glucose levels increase.

We can now derive a formula for estimating the carrying capacity C from lab tests. Assuming that beta cells are at steady state, Eq (4) indicates that C=G B/(G-G0). Estimating B using the well established HOMA-B formula for beta cell function, HOMA-B=20 I/(G-3.5) (with G,I in unit of mmol/L, *µ*U/ml respectively), we arrive at a formula for carrying capacity that we call HOMA-C:

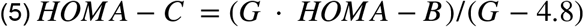

or in terms of insulin and glucose test values (in mM units for G):

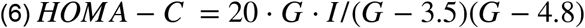

Note that this formula should only be used in cases where glucose is sufficiently higher than 4.8mM, namely in prediabetes or diabetes, in order to avoid small values in the denominator that can increase errors. Unlike HOMA-B which estimates current beta cell function, HOMA-C estimates the maximal beta cell function that an individual can potentially achieve, for example if insulin resistance would rise to very high levels.

### Carrying capacity varies widely across individuals

We used the mathematical model of Eq (1,2,4) to estimate the carrying capacity using data from a cohort of *n* = 256 unmedicated people with prediabetes (17,18). Of these, 8.5% had high enough glucose to be considered diabetic (*G* ≥ 6.9 *m M* /*L*). Each participant had 1-4 measurements over one year of both insulin and glucose.

We considered the tests performed upon enrollment in the study. The distribution of glucose and insulin measurements are widely spread (Fig. 1A). This can be explained by our model with differing values of carrying capacity for each individual (the curves in Fig. 1A). Individuals with low carrying capacity fail to maintain high insulin levels required for glucose homeostasis at high insulin resistance, in contrast to individuals with high carrying capacity.

**Figure 1.**
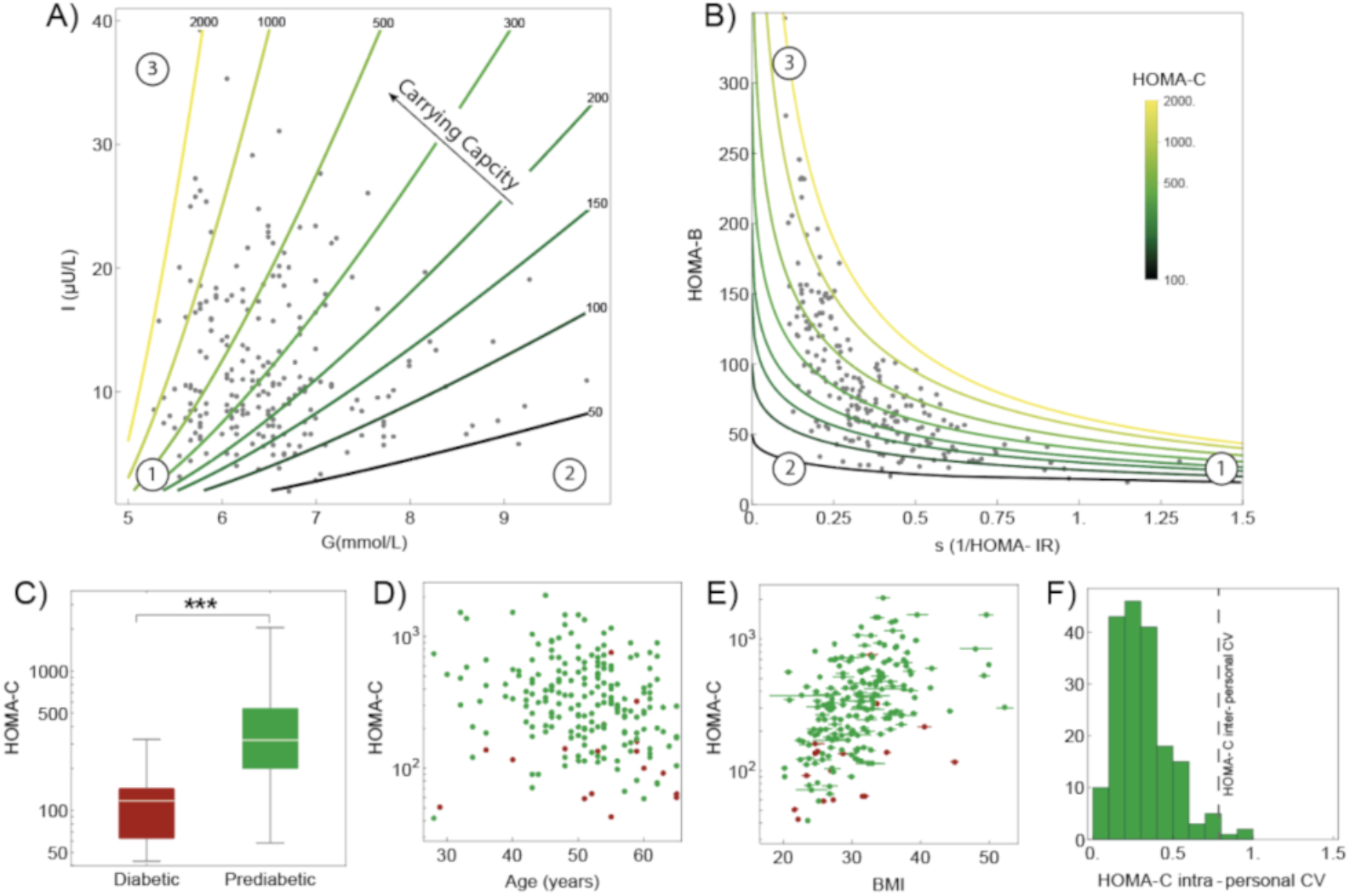
Carrying capacity estimated from insulin and glucose measurements from 256 unmedicated participants with prediabetes from (17,18). (A) Insulin and glucose in the unmedicated prediabetes cohort at enrollment, with outliers removed by excluding data outside of the 90% density contour. Curves denote different values of carrying capacity HOMA-C. (B) Beta cell funcHon (HOMA-B) versus insulin sensiHvity (1/HOMA-IR). Curves show different values of carrying capacity HOMA-C. Numbered circles are equivalent points in A and B for orientaHon. (C) HOMA-C in individuals with prediabetes and diabetes according to the criteria of(17). (D) HOMA-C versus parHcipant age. (E) HOMA-C versus parHcipant BMI. (F) DistribuHon of the coefficient of variaHon (CV) of HOMA-C within an individual’s measurements. VerHcal line shows the CV of the mean HOMA-C between individuals. Formulae are HOMA-B=20 I/(G-3.5), HOMA-IR=I G /22.5 and HOMA-C=G HOMA-B/(G-4.8).

The data can also be visualized in terms of insulin sensitivity and beta cell function (Fig. 1B). As insulin resistance rises, beta cell mass rises and asymptotically reaches the carrying capacity when *Si* = 0.

Figure 1A shows that the predicted carrying capacity HOMA-C varies by about 10-fold between individuals. This relates to postmortem histology studies that report a 5-fold variation in total beta cell mass between equal aged individuals (28).

We compared the carrying capacity between individuals with prediabetes and those with diabetes. HOMA-C was 2.5-fold lower on average in those with diabetes (Fig. 1C, *p*=3 10^-7^). This is in line with postmortem histology studies that showed 50% lower beta cell total mass on average in individuals with diabetes (13).

The HOMA-C carrying capacity does not vary significantly with participant age (Fig. 1D). This indicates that age as a risk factor for diabetes might not operate by reducing carrying capacity, but via other mechanisms such as increased insulin resistance.

Carrying capacity shows mild variation with BMI, rising by a factor of about 2 on average across the range of BMI in the cohort (Fig. 1E). This may be due to the cohort composition which includes primarily individuals with prediabetes and not full-blown diabetes. Individuals with high BMI tend to have high insulin resistance and thus transition to diabetes rather than remain prediabetic unless they have a high carrying capacity.

Repeating this analysis using all timepoints in the dataset shows essentially the same conclusions (Fig. S2). We further analyzed a larger dataset containing 20 years of blood tests (that is 7624 glucose and insulin measurements from Centers for Disease Control and Prevention, National Health and Nutrition Examination Survey (29)). Similarly HOMA-C was found to be almost 3-fold larger in the prediabetic group than is the diabetic one. It varied weakly with age and only moderately with patients’ BMI (Fig. S3)

We next asked how the predicted carrying capacity changes in each individual over time. We computed the variation of HOMA-C between the longitudinal measurements of each individual over a year (Fig. 1F). This intrapersonal variation is about 25% on average, as quantified by the coefficient of variation (CV=standard deviation/mean). In contrast, HOMA-C varies between individuals by about CV=75% (Fig. 1D, vertical line). Thus, carrying capacity in this dataset may be regarded as a relatively constant trait of an individual.

To test whether low HOMA-C is a risk factor for transitioning from prediabetes to diabetes in this cohort, we considered participants with prediabetes (G<6.9mM) at enrollment, evaluated their HOMA-C at enrollment, and asked which participants transit to diabetes over the year of followup. There were 188 participants with at least one followup glucose test, and the criterion for transition was a glucose test above 6.9mM. A total of 44 participants transitioned to diabetes. The risk ratio between the lowest and highest quartiles of HOMA-C was 4+/-0.24 (*p*=0.003). The risk ratio per quartile of HOMA-C was 1.7+/-0.6 (p=0.0007). We conclude that low carrying capacity is a risk factor to transition from prediabetes to diabetes in this cohort, consistent with the notion that low carrying capacity reduces the ability of beta cells to compensate.

### Insulin resistance changes with a timescale of about one year in prediabetes

So far, we have introduced the notion of beta cell carrying capacity within a mathematical model for diabetes, deduced a simple HOMA-like formula for it, and analyzed its values for unmedicated prediabetic patients.

To better understand the dynamics of transition to prediabetes and diabetes, we need to calibrate the timescales in the model. We focus on two timescales: the rate of change of insulin resistance in prediabetes, and the rate of compensation by beta cells.

We first estimated the rate of change of insulin resistance in prediabetes. For this purpose we analyze the longitudinal data from the cohort of *n* = 256 unmedicated people with prediabetes used above (17,18). We used these data to estimate the rate of change of insulin resistance, defined as the inverse of insulin sensitivity, *1/Si*, calculated using HOMA-IR (7). To define the rate of change of *Si* we fit exponential dynamics, *Si*~*e*^*αt*^, to every two consecutive measurements. Such exponential dynamics were proposed by Topp et al. (9), but any rising dynamics with a characteristic timescale would suffice to assess the rate of change of insulin resistance. We therefore defined the rate of change of insulin sensitivity between every two consecutive measurements as 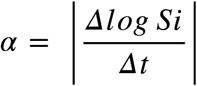 where *α* has units of 1/time. This yields a timescale for changes in insulin resistance of *T*_*R*_ = 1/*α*.

The timescale for insulin resistance collected from all consecutive pairs of measurements has a broad distribution (Fig. 2). The mode is at *T*_*R*_= 362 ± 41 days and 95% of the data exceeds 104 days. The timescale is similar for males and females (*T*_*R*_ = 375 ± 62 days for males, *T*_*R*_= 340 ± 35 days for women). The timescale for individuals with diabetes is also similar (*T* _*R*_= 344 ± 103 days), as it is for upper and lower quartiles of BMI and age (*T*_*R*_= 282 ± 70 *and* 346 ± 83; 314 ± 41 *and* 343 ± 50 days) respectively. We conclude that the timescale of changes in insulin resistance is about one year in the prediabetes cohort.

**Figure 2.**
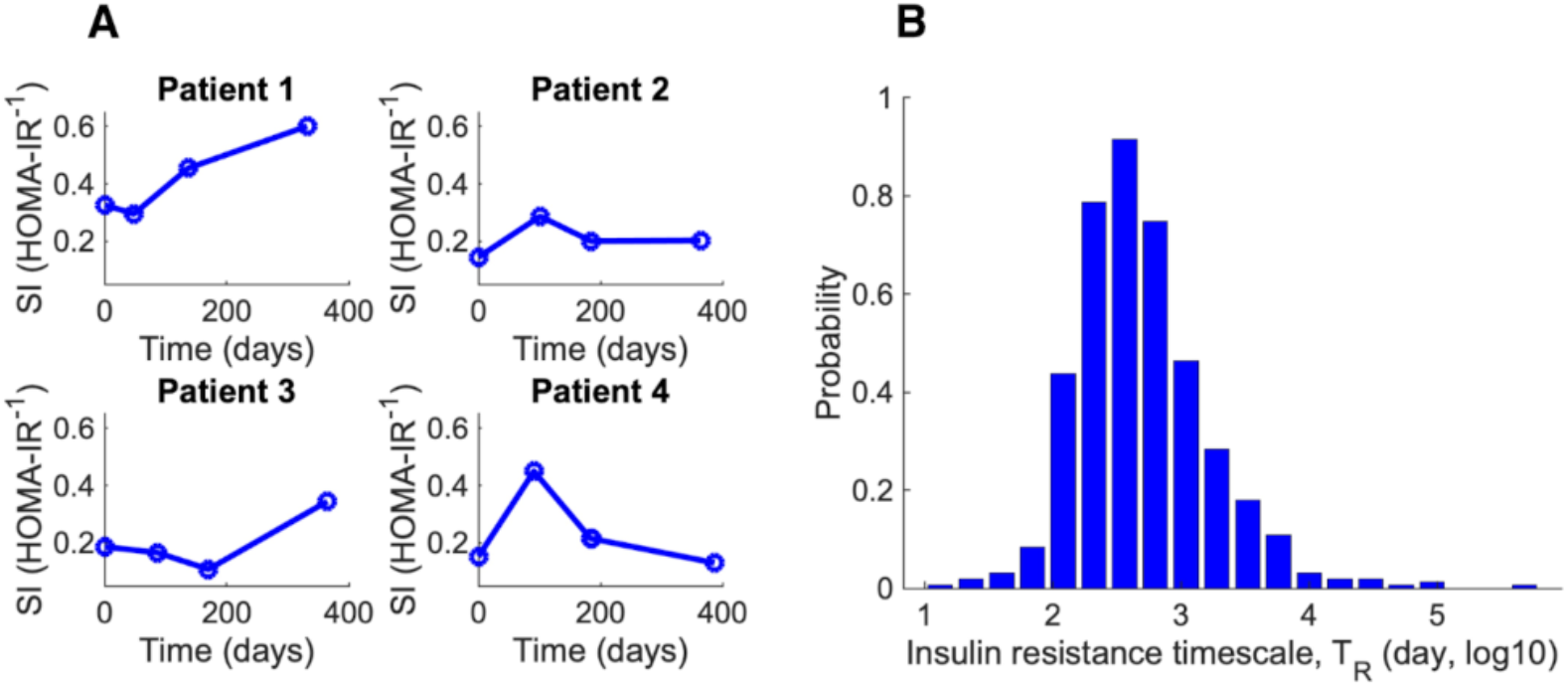
Slow changes in insulin resistance on the scale of a year estimated from a prediabetic cohort. (A) Dynamics of insulin sensitivity (defined as 1/HOMA-IR) for 4 patients over time. (B) Histogram of the timescale of changes in insulin resistance for all pairs of consecutive measurements in the cohort.

### Beta-cell compensation changes on the scale of a month

We next assessed the rate at which beta cells can compensate for changes in insulin sensitivity. This is captured by the rate of change of beta cell functional mass in the model, the parameter *µ* in Eq 4. This describes upregulation of insulin secretion capability per unit beta cell biomass (11) and an increase in total beta cell biomass (9,10), primarily by hypertrophy in humans.

To estimate the compensation timescale, we used two approaches. In both approaches we study glucose when physiology is exposed to changes and observe how rapidly glucose levels adapt after the change. The first approach is based on postpartum recovery of glucose following delivery. During the last two trimesters of pregnancy, insulin resistance rises due to the production of placental hormones (30). With delivery, the placenta leaves the mothers body and insulin resistance becomes restored to its pre-pregnancy value within 2 to 3 days (31). This is an example of the rapid changes in insulin resistance that occur in physiological situations.

We analyzed glucose lab tests after delivery using an electronic medical record dataset from a large Israeli health service, Clalit (32,33) which includes 231260 glucose tests performed in the 20 months after delivery. Mean glucose drops below baseline after delivery by about 2%, a change that is accurately detected due to the large number of glucose tests in this dataset (Fig. 3A). To estimate the rate of compensation, we fit an exponential function *G* (*t*) = *G*_*st*_ − *Ae*^−*λt*^ and defined the timescale for compensation as *T*_*comp*_= 1/*λ*. Glucose recovers with a timescale of *T*_*comp*_=109 +/- 7 days (Fig. 3A). The detailed analysis is given in the Methods section.

**Figure 3.**
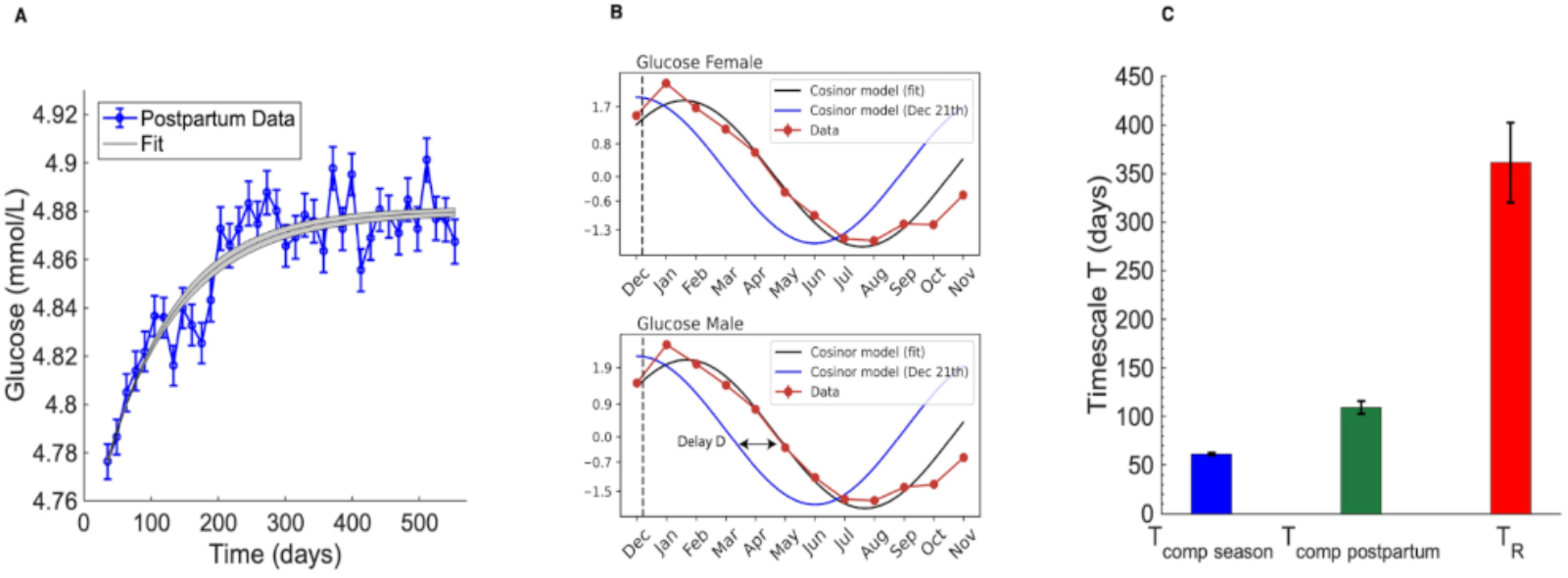
Beta cell compensation is estimated to be about 3-6 times faster than changes in insulin resistance during prediabetes. (A) Mean of glucose blood tests averaged over each week after delivery from the Clalit dataset (total of n=174,344 tests). The beta cell compensation timescale was estimated using an exponential fit. (B) Seasonal variation in mean fasting glucose blood test from the Clalit dataset (in red) along with a fit to a phase shifted cosine (gray line). Normalized photoperiod input, peaking at Dec 21, is shown in blue. Total number of tests is n=5,221,865 (females) and n=3,752,163 (males). (C) Comparison of the beta-cell compensation timescale from seasonality and postpartum estimates and the insulin resistance timescale from the prediabetes cohort. For seasonality and insulin resistance the mean of male and female timescales is shown.

The second approach to estimate the compensation timescale uses seasonality. Like most blood chemistry variables, glucose lab tests show seasonal variations in humans (34). A recent large-scale analysis examined seasonality by means of 9 million glucose tests from the Clalit dataset which includes 50 million life-years (34) (Fig. 3B). This study controlled for circadian effects and excluded data from participants with medical conditions and drugs that affect the glucose test. Blood glucose showed seasonal oscillations with an amplitude of about 2% (Fig. 3B).

For our purpose, the important variable in the seasonality is the delay of the glucose seasonal oscillations relative to the input signal, photoperiod (day length) variation. A system with immediate compensation would lock to the photoperiod signal and show a maximum at December 21, the shortest day, and a minimum at June 21, the longest day. Instead, the seasonal oscillations of glucose tests show a delay relative to the seasonal variation in photoperiod. We estimate this delay by fitting a cosine function with a phase delay to the seasonal variation of glucose tests. This provides a delay of 45 +/- 0.5 days for women and 50 +/- 0.5 days for men (Fig. 3B), leading to *T*_*comp*_=61 +/- 1 days (see Methods for details). We repeated the seasonality delay analysis when binning individuals by age. The estimated compensation time does not appreciably change with age (Fig. S1).

Both approaches, postpartum recovery and seasonality, yield comparable estimates for the beta-cell compensation timescale, *T*_*comp*_ ~ 50-100 days (detailed analysis given in the Methods section).

We conclude that beta cells can compensate at least 3 times faster than insulin resistance typically changes in prediabetes.

### Prediabetes is not consistent with mathematical models without carrying capacity

In this section we suggest that the classic Topp model for beta cell compensation (which lacks a carrying capacity) (9) cannot explain prediabetes given the present estimates in which compensation is faster than the changes of insulin resistance in prediabetes. The intuitive reason is that beta cells can easily compensate for the slow changes in insulin resistance because of their faster timescale. As a result, previous models predict that glucose should not deviate much from its baseline even if insulin resistance reaches high levels - beta cells are always able to catch up and compensate.

The principal dynamical model for compensation is based on the pioneering work of Topp et al. (9), with extensions by Ha et al. (11) and Karin et al. (10). Topp et al. proposed three pathways to diabetes. Two are static, with hyperglycemia causing glucotoxicity of beta cells. These mechanisms are not relevant to prediabetes because they do not explain how glucose can slowly rise from normal levels over years. Instead they predict an abrupt jump from normal to high glucose levels.

The third pathway proposed by Topp et al. (9) is relevant to prediabetes. It is called dynamic hyperglycemia, and occurs when insulin resistance changes faster than beta cells can catch up. We use the Topp model, and show that prediabetes, namely lack of compensation, requires that insulin resistance changes as fast as or faster than beta cell compensation (Fig. 4A). In humans, however, our estimates show that insulin resistance changes 3-6 times slower than compensation. Thus the dynamic hyperglycemia mechanism cannot explain prediabetes. In the Topp model, the rapid compensation that we observe would lead to glucose levels that remain close to baseline (Fig. 4A).

**Figure 4.**
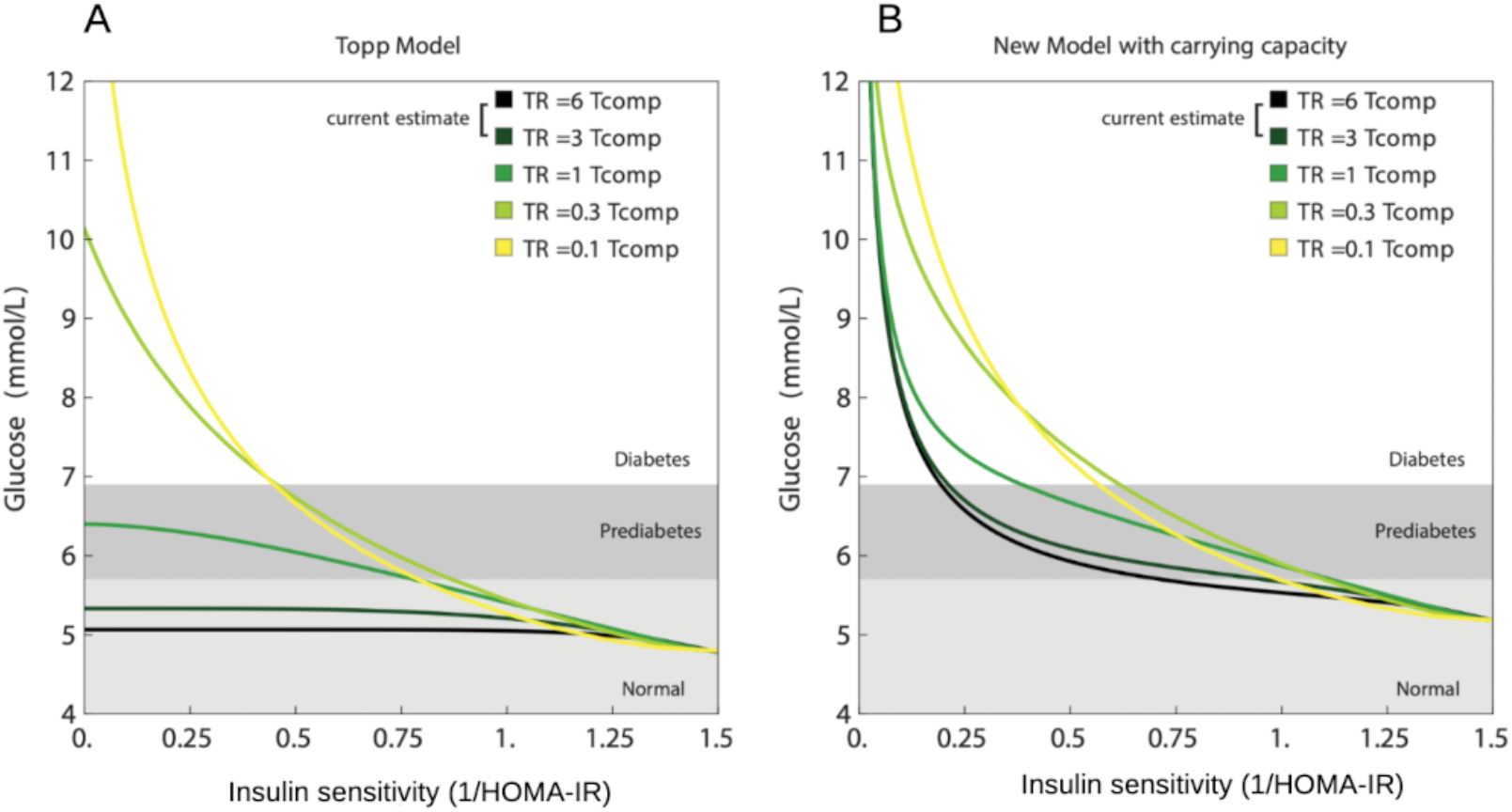
New model with carrying capacity can explain prediabetes in contrast to the Topp model which has no carrying capacity. (A) A model with no carrying capacity, the Topp model, shows glucose near baseline when the compensation time is 3-6 times slower than the rate of change of insulin resistance (black and dark blue curves). (B) New model with beta cell carrying capacity shows prediabetes at all time scale ratios, including the estimated ones. Kinetic parameters:,, and for both models (9), and for the carrying capacity model and. Simulations used exponentially rising insulin resistance with timescale starting from Si =1.5.

The dynamic hyperglycemia mechanism might apply to situations with rapidly increasing insulin resistance which generates higher glucose than baseline. For example, pregnancy, infection and acute stress all cause rapid changes in insulin resistance and can show glucose levels that differ from baseline. Indeed, this is presumably the physiological role of rapid changes in *Si* caused by these conditions- to direct glucose to the fetus, immune system and brain, respectively, at the expense of muscle and fat cells (35). But for prediabetes - in which glucose levels rise over years - a different etiology is required.

The present carrying capacity model shows the slow rise of glucose even with the relatively rapid compensation observed here Fig 4B. We conclude that given the slow rise in insulin resistance in prediabetes, the present model can explain prediabetes whereas previous models cannot.

### Integrated model for prediabetes and diabetes subtypes

The carrying capacity model provides an integrated picture of beta cell function and glucose levels that describes the transitions between normoglycemic, prediabetic and diabetic conditions. In addition, it mathematically describes different diabetes subtypes.

To provide intuition, we use a graphic approach to understand the predictions of the model. This graphic approach, called nullcline analysis, is widely used in the study of dynamical systems (36). It highlights the predictions of the model based on its two feedback arms (Fig. 5A). One arm describes the decrease in steady-state glucose due to beta cell function: the higher beta cell function the lower steady-state glucose due to insulin. This is called the *G*-nullcline. The other arm is the increase in steady-state beta cell functional mass *B* due to glucose, which underlies compensation. This is called the *B*-nullcline. The carrying capacity is seen in the shape of the *B*-nullcline, which saturates at a maximum value at the carrying capacity. These two nullclines cross at a point which defines the system’s fixed point, because it is the point where both *G* and *B* are at steady state. The fixed point determines the steady-state glucose and beta cell function mass.

**Figure 5.**
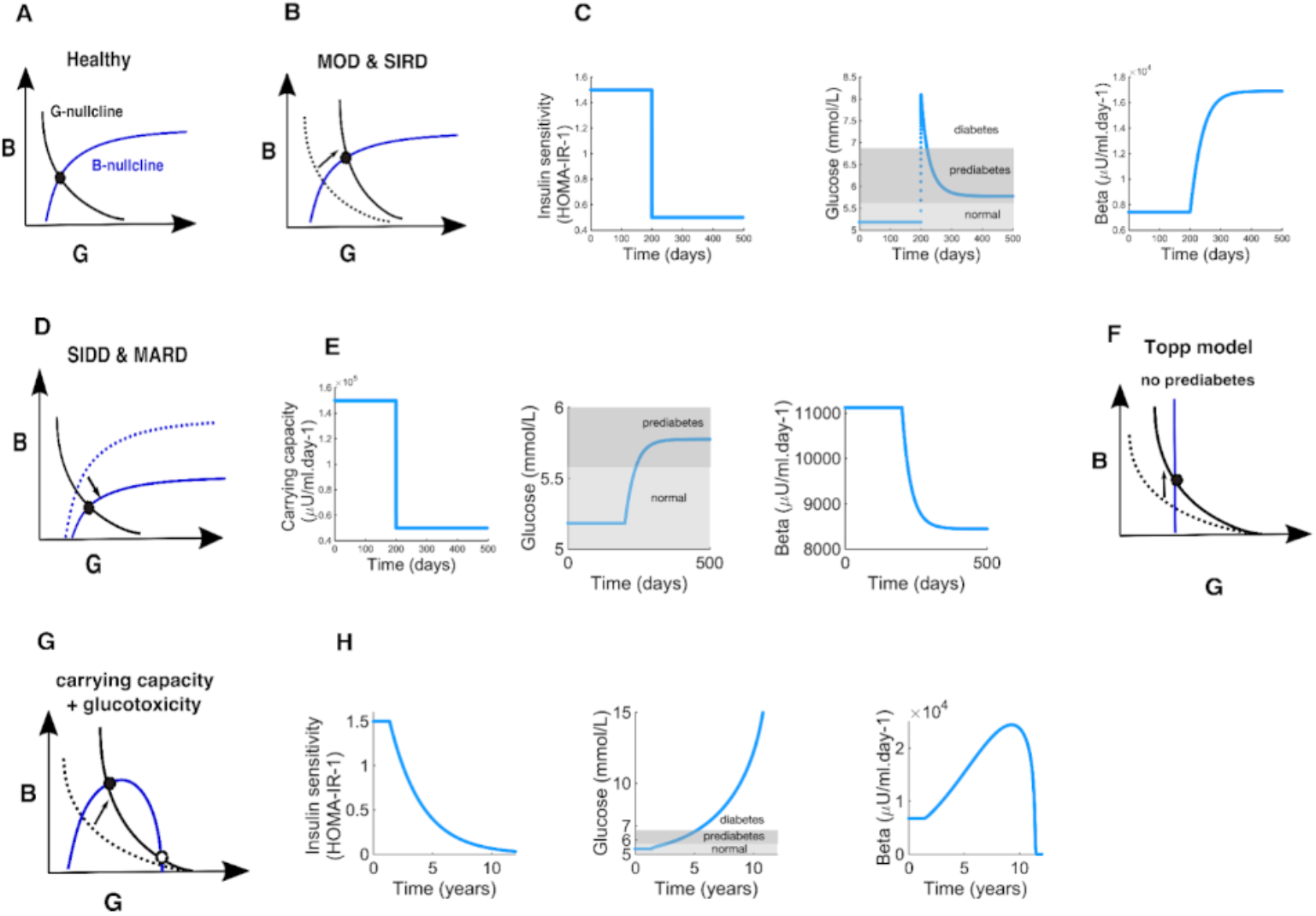
The carrying capacity model provides a framework to understand changes in steady state glucose and beta cell functional mass. (A) Nullclines for the carrying capacity model. (B) Effects of raising the G nullcline as in insulin resistance. (C) Dynamics of the carrying capacity model for a step-like decrease in insulin sensitivity from to. (D) Effects of lowering the -nullcline as a result of a reduction of carrying capacity. (E) Dynamics of the carrying capacity model for a step-like decrease in the carrying capacity. The carrying capacity drops from to. (F) Nullclines for the Topp model. (G) Nullclines for the model with carrying capacity and glucotoxicity. (H) dynamics of the glucotoxicity model for an exponential increase in insulin resistance with. Kinetic parameters:,,,, and G1=10 mmol/L.

Physiological conditions and medical interventions can change these nullclines, thus shifting the fixed point and hence the steady-state value of glucose and beta cell functional mass. Some physiological conditions affect the *G*-nullcline, namely the steady state glucose at a given beta cell mass (Fig. 5B). This includes insulin resistance, which shifts the nullcline to higher levels of glucose, due to the reduced effectiveness of insulin. This causes a rise in both steady state glucose and beta cell functional mass. Other conditions that affect the *G*-nullcline are excess glucose production by the liver, excess insulin removal rate, reduced insulin-independent glucose removal, and impairments in the beta-cell response to glucose. All of these conditions in the model show a rise in beta cell mass together with a rise in glucose. For example, upon a step-like rise in insulin resistance, the model shows how glucose rises within hours, and then declines more slowly over weeks as beta cell mass increases (Fig. 5C).

In contrast, other conditions affect the *B*-nullcline. This includes a reduction in beta cell health, so that a given glucose level results in lower beta cell functional mass pushing the -nullcline to lower values (Fig. 5D). Such changes also shift the fixed point. They give rise to higher glucose levels, but lower beta cell function mass. A similar effect is caused by conditions that lower the beta cell carrying capacity.

Thus changes in the *G*-nullcline should show increases in the the steady state values of both *G* and *B*, whereas changes in the *B*-nullcline should show an increase of but a decline of *G*. but a decline of *B* Such a drop in beta cell secretion capability causes a rapid rise in glucose followed by a decline as beta cell mass grows to compensate (Fig. 5C).

Using the distinction between the two nullclines, one can analyze different subtypes of diabetes. Recently, four prevalent subtypes of non-autoimmune diabetes were proposed (37). Two of these, mild obesity related diabetes (MOD) and severe insulin resistance diabetes (SIRD), are accompanied by a rise of insulin resistance, corresponding to an increase in the *G*-nullcline. Indeed these types show a rise in both *G* and beta cell function *B* (Fig. 5B).

The other two types, severe insulin deficient diabetes (SIDD) and moderate age-related diabetes (MARD) have the opposite phenotype- a rise in glucose but a decline in beta cell functional mass (37). We propose that these two types of diabetes involve a shift in the *B*-nullcline, compromising beta cell compensation rather than insulin resistance as a primary etiology (Fig. 5D).

A similar analysis can be made for mature onset diabetes of the young (MODY), which accounts for about 5% of diabetes cases. Most MODY cases are due to two major genetic variations. The HNF1 variants (50% of MODY) impair insulin secretion, lowering the *B*-nullcline and indeed show the predicted reduced B-cell function along with high glucose (38). The other variant, GCK (30%), impairs glucose sensing in beta cells. It thus should affect both nullclines (see nullcline equations in Methods). It has higher beta-cell function than HNF1 (3-fold) and non-MODY T2D (1.5 fold) (38).

Notably, the Topp model without carrying capacity has different nullclines that cannot capture these subtypes of diabetes (Fig. 5F). The Topp model has the same -nullcline as the present model, but the *B*-nullcline is a vertical line at *G* = *G*_0_. At steady state, compensation always brings glucose back to baseline. The present model differs from the Topp model by having a carrying capacity, and this is the origin of the curved shape of the *B*-nullcline and the existence of prediabetes steady states with glucose above baseline (Fig. 5B,D).

Finally, we added glucotoxicity to the carrying-capacity model (see SI). Glucotoxicity is an effect, characterized primarily in rodents, in which beta cells are removed or become dysfunctional when glucose reaches very high levels (39,9,16,40). Glucotoxicity causes the B-nullcline to have a rising and falling shape, since beta cell mass shrinks at high glucose levels (Fig. 5G). The two nullclines now cross at three points: a stable fixed point at normal glucose levels and another a stable fixed point at very high glucose in which beta cells mass is zero. Between these is an unstable fixed point.

Rising insulin resistance pushes the *G*-nullcline up, and moves the low stable fixed point and the unstable fixed points closer to each other (Fig. 5G). Glucose baseline thus rises as in prediabetes and into the range of type-2 diabetes. At a critical level of insulin resistance the system goes through a saddle-node bifurcation at which the two fixed points collide and annihilate, leaving the high glucose fixed point as the only steady-state solution. Such a process may correspond to a transition to insulin-dependent diabetes at very high insulin resistance (Fig. 5H).

## Discussion

We have presented a mathematical model for the carrying capacity of beta cell compensation. We have provided a HOMA-C formula to estimate the beta cell carrying capacity in an individual - the predicted maximal beta cell function achievable-based on glucose and insulin blood tests. Using a cohort of unmedicated people with prediabetes, we found that inferred carrying capacity varies by up to 10-fold between individuals. The lowest quartile of carrying capacity was associated with a 4-fold higher risk to develop diabetes compared to the top quartile.

We found that the rate of change of insulin resistance in participants with prediabetes has a typical timescale of about a year. We also estimated the timescale for beta cell functional compensation using large medical datasets on glucose postpartum recovery and seasonality. Both estimates point to a compensation timescale of 50-100 days, about 3-6 times faster than the timescale of changes in insulin resistance during prediabetes. We concluded that beta cell functional mass can easily track and compensate for insulin resistance changes in prediabetes, and hence existing mathematical models of compensation cannot explain the slow rise of glucose in prediabetes. The carrying capacity of beta cell function can explain prediabetes - compensation reaches a limit when beta cells approach their carrying capacity; further rise in insulin resistance causes glucose to rise. The new model shows prediabetes even when insulin resistance rises very slowly.

Slow changes in insulin sensitivity in prediabetes, on the scale of a year, are consistent with findings from recently diagnosed patients with T2D from the Innovative Medicines Initiative Diabetes Research on Patient Stratification (IMI-DIRECT) study (41). Insulin sensitivity decreased by a factor of about 2 over 36 months in a subpopulation of fast progressors, and showed no change on average in the remainder 90% of the population. Similarly, slow rates of change of HOMA-IR in individuals with risk of type-2 diabetes were documented in the Botnia study (42).

The concept of a carrying capacity for beta cell compensation requires further research. It would be important to study what sets the beta-cell carrying capacity, and how it differs between individuals and across physiological conditions. Carrying capacity may be affected by conditions that modulate maximal insulin secretion per beta cell unit mass. These include incretin hormones like GLP1, neuronal inputs to beta cells and drugs like sulfourea, all of which increase beta cell insulin release, and somatostatin secreted by delta cells which reduces insulin release.

Reduced carrying capacity is predicted to be a major risk factor for prediabetes and diabetes in this mathematical model. It would be interesting to study whether and how carrying capacity declines with conditions that raise the risk of diabetes. A recent review summarizes evidence that total beta cell mass and function is lower in prediabetes and in T2D patients on the order of 20-50% compared to healthy individuals (13).

One relevant suggestion is that the total number of beta cells, determined during development and childhood, varies between people and affects the risk for diabetes in adulthood (20). The 5-fold variation in beta cell total mass between equal-aged individuals (28) may underlie some of the 10-fold variation in carrying capacity estimated here within the unmedicated prediabetes cohort.

Our model provides a way to estimate carrying capacity from glucose and insulin measurements. We call this the HOMA-C equation, in analogy to the widely used HOMA-IR and HOMA-B equations, where HOMA stands for homeostatic model assessment (7). The HOMA-C equation is HOMA-C=G HOMA-B/(G-4.8), with HOMA-B=20 I/(G-3.5) where *I* is fasting insulin in units of U/ml and *G* is fasting glucose in units of mmol/L. HOMA-C can only be used for individuals with glucose above the normal range and in conditions where beta cell functional mass can be assumed to be close to steady state. It should not be used proximally to conditions that cause abrupt changes in insulin resistance such as infections, glucocorticoid treatment or acute stress, because beta cell functional mass compensation has not reached steady-state. This equation may be useful to predict the risk of T2D, where the lower HOMA-C the higher the predicted risk.

Carrying capacity might also be important when insulin resistance rises at a time when other physiological conditions already call for increased beta cell functional mass. For example, during pregnancy, beta cell insulin secretion increases to compensate for pregnancy-induced insulin resistance. An additional rise in insulin resistance due to, say, chronic stress or obesity, would be especially harmful because beta cells are already pushed closer to their carrying capacity by the pregnancy. This may explain part of the risk for gestational diabetes. More generally, carrying capacity can help to explain how different conditions can combine to cause prediabetes and diabetes, by pushing beta cells closer to their carrying capacity.

The concept of a carrying capacity for an endocrine gland has recently been used in a mathematical model in another subclinical endocrinological condition-subclinical hypothyroidism. The thyroid axis glands can change their functional mass to compensate for changes in conditions (10). The thyroid mass famously grows at low iodine, causing Goiter. Kohanim et al. showed that a carrying capacity of the pituitary cells that secrete thyroid stimulating hormone (TSH) can explain subclinical hypothyroidism in a way that is similar to the present mechanism for prediabetes (26). When pituitary cells approach carrying capacity they can no longer secrete additional TSH to compensate for autoimmune killing of the thyroid in Hashimoto’s thyroiditis-causing a transition from subclinical disease to clinical disease with overt hypothyroidism. Similar carrying capacity effects might be at play in other hormonal systems.

The present estimate for the beta cell compensation time *T*_*comp*_ and the timescale for insulin presence change in prediabetes patients *T*_*R*_ can guide the timescale of interventions aimed at reducing glucose. One may use and extend the present HOMA-C formula to understand the carrying capacity of each individual based on glucose and insulin measurements. This may allow clinicians to track treatments of prediabetes in terms of how the treatment moves beta cells away from their carrying capacity.

The present model is consistent with longitudinal studies that showed that before the onset of prediabetes, glucose rises slowly within the normal range over decades (43). Once it crosses about 6mM glucose rises more rapidly over several years until diabetes occurs (43).

Mathematical models have historically been important in clinical research on diabetes (44,45,46,7). The new carrying capacity model and the HOMA-C formula which can be estimated from glucose and insulin lab tests may thus help form a more accurate understanding of prediabetes and its transition to diabetes in each individual.

## Supplementary Materials and Methods

### Carrying capacity model

The Topp model (1) describes the rates of change of blood glucose *G*, insulin *I* and beta cell functional mass *B*, adapted here as follows:

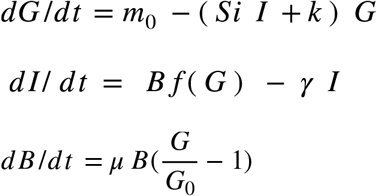

where *m*_0_ is liver and postprandial glucose input, *Si* is insulin sensitivity, *k* is the insulin-independent glucose removal rate. Insulin is secreted by beta cells with mass beta cell mass *B* (in units of insulin concentration per unit time) and removed at rate *γ*. The regulation function of insulin by glucose is a sigmoidal function 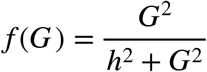 with halfway point *h*. The beta cell functional mass changes with growth rate *µ* to reach the glucose set point *G*_0_.

We ignored the glucotoxicity term at very high glucose levels in the original model 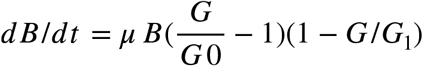. Ignoring this (1 − *G /G*_1_) term does not change the conclusions of this study, because the glucose levels in the present study are well below the glucotoxicity threshold of *G*_1_= 23 mmol/L in Topp et al. (1). This system can be simplified to a set of 2 ordinary differential equations (ODEs) by means of separation of timescales. Insulin is removed with a half life of about 5 minutes which is faster than glucose removal on the scale of hours and beta cell functional mass growth on the scale of weeks. Insulin thus equilibrates rapidly compared to the slow timescale, so that we use a quasi steady state assumption with *dI* / *dt* = 0. The quasi-steady-state solution is *I*_*qss*_ = *q B f* (*G*) *γ* ^−1^. Substituting this expression for *I* in the equation for glucose reduces the system to two equations:

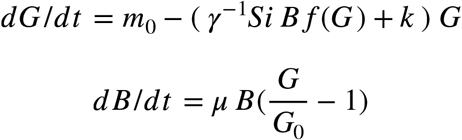

We use the kinetic parameter values used by (1,2), (Table S1.).

The new model extends the Topp model by introducing a carrying capacity for beta cells, *C*. Introducing the latter in the 2 ODE version of the Topp model, we obtain

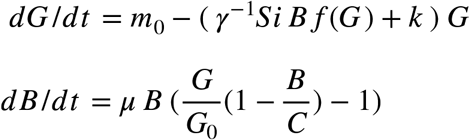

where *C* is the carrying capacity.

In the limit of very slow changes in *Si* relative to the compensation time, which is appropriate to the present estimates, one obtains the following relation between glucose and insulin sensitivity *Si*

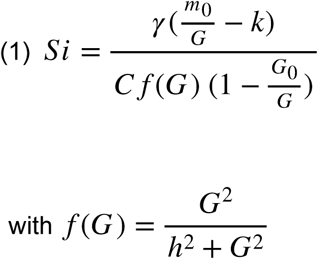

The relation between functional beta cell mass *B* and *Si* is

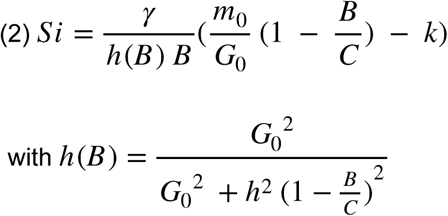

Based on this model, we define the HOMA-C formula based on the steady state assumption for *B*, namely *dB/dt=0*. This provides *C* = *G B* /(*G* − *G*_0_). Since B is widely estimated using the HOMA-B formula, HOMA-B=20 I/(G-3.5), we propose the following formula for carrying capacity 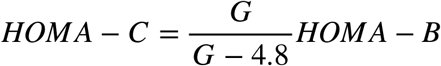 with *G*_0_ =4.8 mmol/L.

### Data from an unmedicated prediabetic cohort

We analyze data from a cohort described in (3,4). The cohort consists of unmedicated prediabetic and diabetic adults who underwent a 6-month dietary intervention and additional 6-month follow-up with several (up to four) blood tests of fasting glucose and insulin levels. We only analyzed data from patients with at least one fasting glucose and insulin measure simultaneously, leading to 256 participants (111 women aged 50 ± 8 *years*, 145 men aged 51 ± 7 *years*), 8.5% being diabetic at the beginning of the dietary intervention. Patients were included in the diabetic cohort if either their blood glucose or their HbA1c levels were elevated (5).

Insulin was measured with the use of a chemiluminescent microparticle immunoassay (ARCHITECT insulin assay) with precision of less than 7% CV and plasma glucose was measured with the use of a hexokinase method (GLUC2 assay, cobas; Roche) with precision of about 2% CV.

From these data, we extract how individual insulin sensitivities and functional beta cell mass evolve over time using the HOMA-IR and HOMA-B indices (5), *HOMA − IR = G I*/22.5 and *HOMA* − *B* = 20 *I* /(*G* − 3.5) where *G* and *I* are fasting glucose in mmol/L and insulin in *µ*U/ml.

As in (1), we defined the rate of change of insulin sensitivity as *α =* |*1/Si dSi/dt*|. We calculated the rate of change between each measurement for each patient of the cohort as 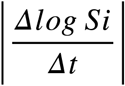.

### Data from the National Health and Nutrition Examination Survey (CDC)

We analyze data from the National Health and Nutrition Examination Survey (CDC) (6). It consists of blood tests (fasting glucose and insulin) from 1999 to 2018. After filtering out the outliers for glucose, insulin and BMI, we obtained 7624 data points. Among them are 83% m prediabetic and 16 % diabetic individuals (those who reported taking insulin or glucose lowering medications excluded). For prediabetic individuals, mean HOMA-C= 640.5 +/- 5.6 (mean+/- SEM) and for diabetic ones, mean HOMA-C=215.7 +/- 5. Whether HOMA-C is predictive of the risk to transition from prediabetes to diabetes could not be tested with this cross-sectional dataset because there was no longitudinal follow-up.

### Postpartum glucose data from the Clalit medical record dataset

We analyzed glucose postpartum dynamics using fasting blood glucose tests from the Clalit (a major Israeli health service organization) medical database from women after their first delivery (7). We analyzed anonymized data from two weeks postpartum to 80 weeks after delivery.

### Seasonal glucose data from the Clalit medical record dataset

To analyze glucose seasonality we use data from Tendler et al. (8). Briefly, these data consist of hormone and metabolite levels from the Clalit medical database for healthy individuals from 20 to 80 years. The results that were binned into 12-month bins and the mean percentile value was calculated for each bin.The seasonality of the percentile change was quantified by fitting the data to a cosinor model *y*(*t*) = *A cos*(*ω t* + *𝜙*) + *e*(*t*) where *A* is the amplitude, *𝜙* is the phase and *e* is the error with respect to the cosinor model (8).

### Seasonal phase shift

In a linear system driven by a periodic input, the delay *D* between input and output can be expressed as a function of the kinetic parameter of the system. Here we use the estimated value of the delay *D* between day length variation (input) and glucose (output) to estimate the beta cell mass turnover rate from the Topp model, with the approximations *k* = 0 and 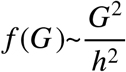. The system has two timescales: after a perturbation, glucose levels equilibrate within hours whereas beta cell mass evolves on a slower timescale. The quasi-steady-state solution for *G* is

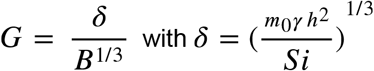

On slow timescales, B cell mass evolves as

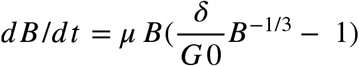

The linearized equation for *B* is

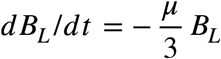

A yearly driving force *f* (*t*) applied on the linearized beta cell mass equation is described by

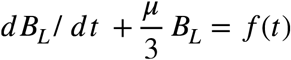

where *f* (*t*) = *sin*(*ω t*) with 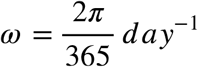.

The transfer function of the system is

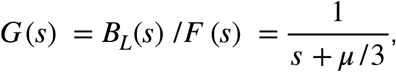

where *B*_*L*_(s) and *F* (*s*) are the Laplace transforms of *B*_*L*_(t) and f(t) respectively. *G* (*s)* can also be expressed as *R exp*(*ϕ*) with gain 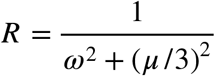 and phase shift is 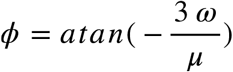 and *ϕ* are *ϕ* the gain and the phase shift between the input f(t) and the output *B*_*L*_(t). Altogether we find

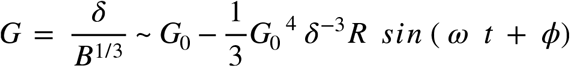

So that the relation between the delay and the beta cell growth rate is

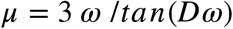

### Postpartum recovery analysis

The analysis of the previous section provides a glucose quasi-steady state of

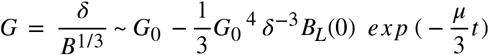

Normalizing the steady state to *G* = 1 and *B* = 1 implies

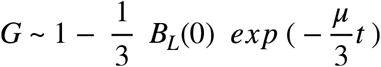

The compensation time can be obtained by fitting a function

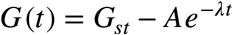

to the data. We get 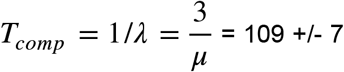 days from the postpartum recovery data. For the seasonal glucose data, we have 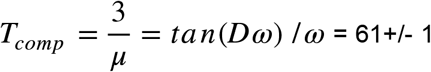 days, which is in the same order of magnitude.

### Carrying capacity model nullclines

We compute the nullclines of the system, that is the curves corresponding to *dG* /*dt =0* and *dB* /*dt =0*. The *G*-nullcline (*dG* /*dt* =0) is

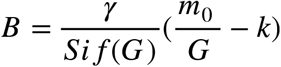

Without carrying capacity (that is, C = ∞), as in the Topp model, the *B*-nullcline is vertical (*G* = *G*_0_) and there is always compensation. However the presence of a finite carrying capacity C bends the *B*-nullcline as

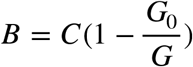

When the value of *Si* decreases, glucose levels remain nearly constant at first because the rise of B cell mass provides compensation. However, when *B* approaches its carrying capacity *C* glucose levels start to rise: this is the onset of prediabetes. Note that when insulin resistance become very high, that is, in the limit of *Si*→0, the state of the system converges to 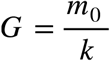 and 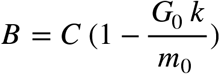.

### Model with carrying capacity and glucotoxicity

The Topp model originally contains a glucotoxicity term (1). If we add such a term to our model with carrying capacity (Fig. 5GH), we obtain

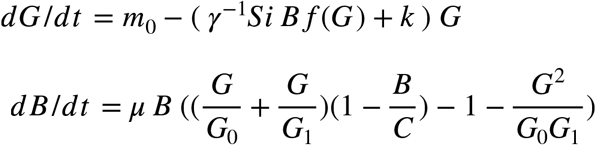

When *B* ≪ *C*, that’s when cells are far from carrying capacity, we recover the original Topp et al equation:

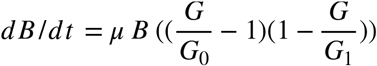

**Fig. S1.**
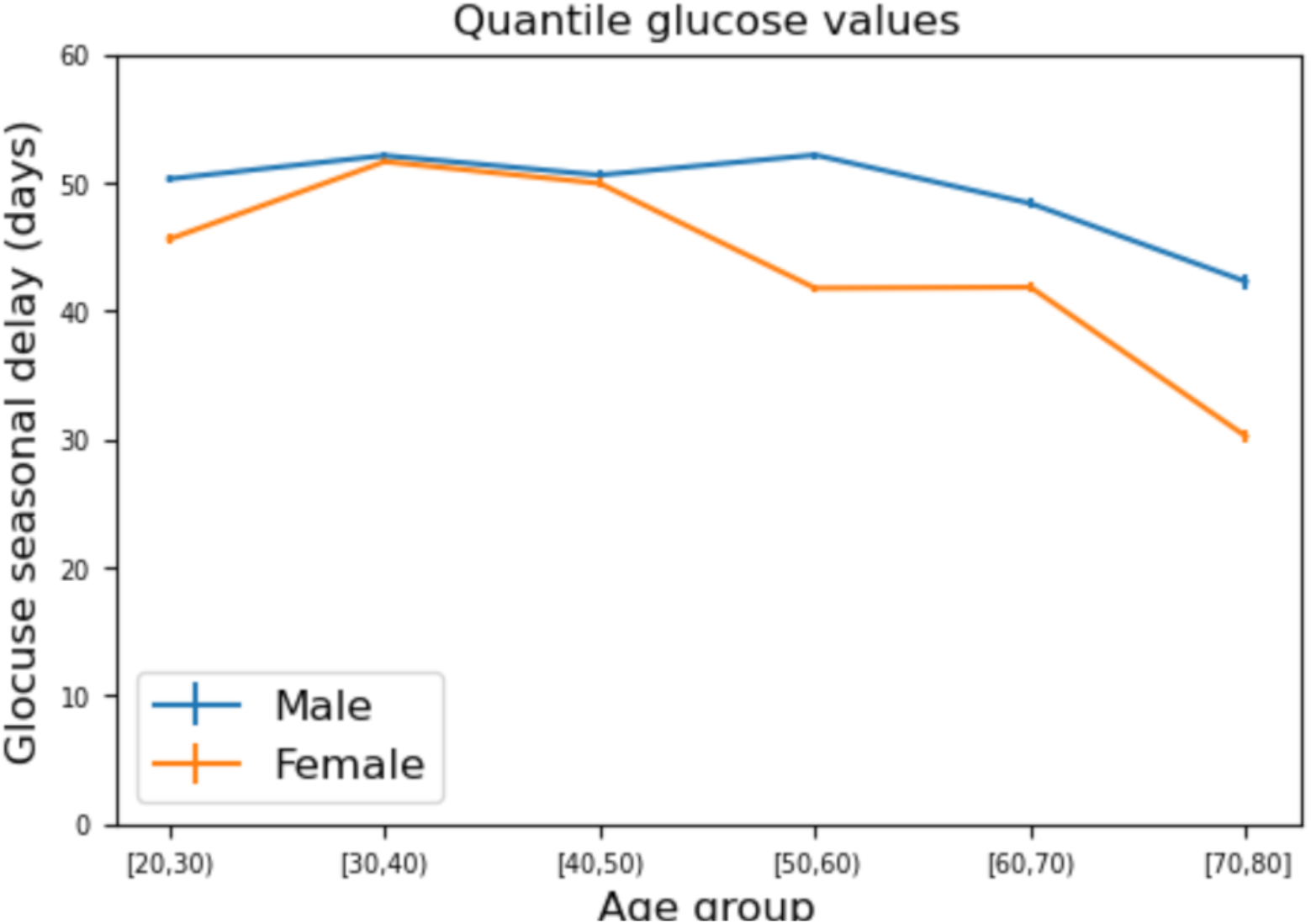
The seasonal delay of glucose lab tests does not increase with age. Seasonal delay was computed for Clalit glucose quantile data binned according to age. Shown are mean and standard error for males and females.

**Fig. S2.**
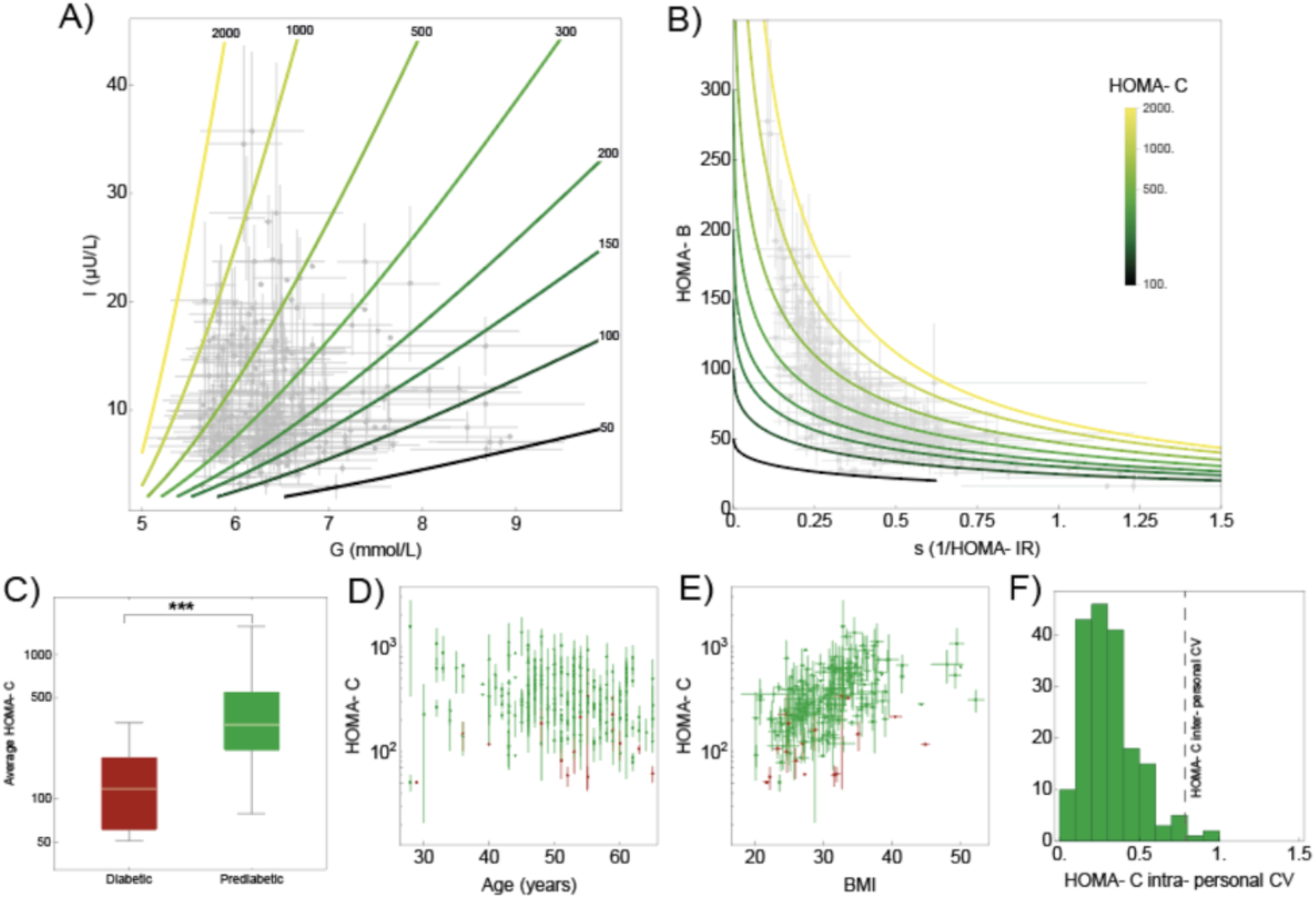
Carrying capacity equation HOMA-C for all insulin and glucose measurements from 256 unmedicated participants with prediabetes from (3,4). Error bars show the std of measurements over all timepoints for each individual. This figure is the same as Fig. 4, except that all measurements are used and not only the first measurement at enrollment.

**Fig. S3.**
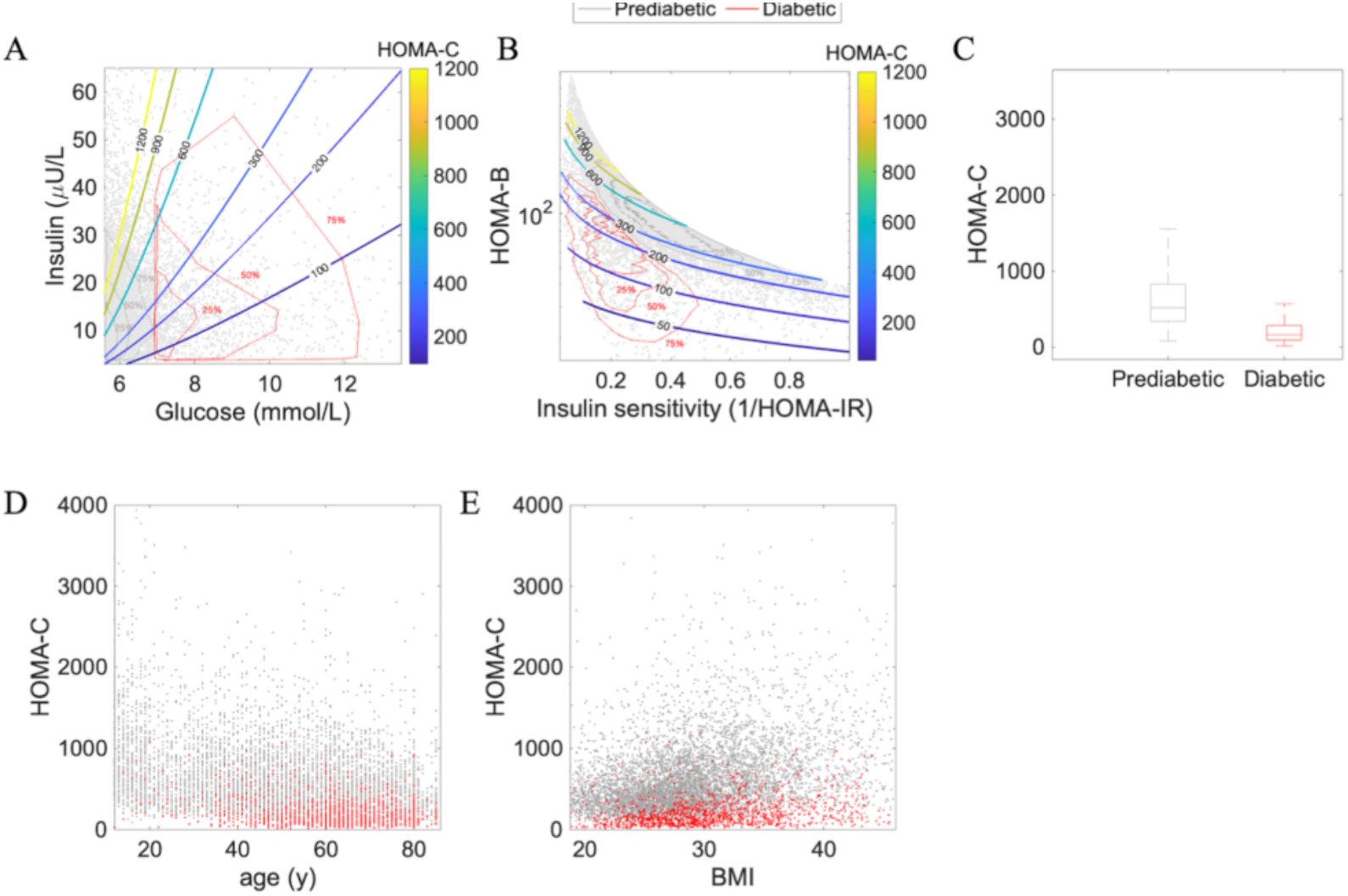
Carrying capacity estimated from insulin and glucose measurements from 7624 unmedicated participants with prediabetes from (6). (A) Insulin and glucose in prediabetic and unmedicated diabetic patients, with outliers removed by excluding data outside of the 95% density contour. Gray contours contain 75%, 50% and 25 % of prediabetic patients respectively. Red contours contain 75%, 50% and 25 % of diabetic patients respectively. Curves denote different values of carrying capacity HOMA-C. (B) Beta cell function (HOMA-B) versus insulin sensitivity (1/HOMA-IR). Curves show different values of carrying capacity HOMA-C. (C) HOMA-C in individuals with prediabetes and diabetes. (D) HOMA-C versus participant age. (E) HOMA-C versus participant BMI. Formulae are HOMA-B=20 I/(G-3.5), HOMA-IR=I G /22.5 and HOMA-C=G HOMA-B/(G-4.8).

## Author Contributions

AW, YT, YoT and UA designed research. AW, YT, AB and AM performed research. AW, YT, AM, AB, MR, AG, NMC, ES and SS analysed data. AW, YT, DG and UA wrote the paper.

## Acknowledgments

UA acknowledges funding by the European Research Council under the European Union’s Horizon 2020 Research and Innovation Programme Grant 856587.

## Code availability

Codes were written in MATLAB_R2022b, in Mathematica 13.2 (Wolfram) and in Python 3.9.7 and can be found in the following link: https://github.com/Erorua/beta_cell_carrying_capacity/blob/main/Code.zip.

